# Discovery of a new species of trichomonasvirus in the human parasite *Trichomonas vaginalis* using transcriptome mining

**DOI:** 10.1101/2021.12.09.471948

**Authors:** Austin R. Manny, Carrie A. Hetzel, Arshan Mizani, Max L. Nibert

## Abstract

*Trichomonas vaginalis* is the most common nonviral cause of sexually transmitted infections globally, with an estimated quarter of a billion people infected around the world. Infection by the protozoan parasite results in the clinical syndrome trichomoniasis, which manifests as an inflammatory syndrome with acute and chronic consequences. Half or more of these parasites are themselves infected with one or more dsRNA viruses which can exacerbate the inflammatory disease. Four distinct viruses have been found in *T. vaginalis* to date, Trichomonas vaginalis virus 1 through 4 (or TVVs). Despite the global prevalence of these viruses, few coding-complete genome sequences have been determined. We conducted viral sequence mining in publicly available transcriptomes across 60 RNA-seq datasets representing 13 distinct *T. vaginalis* isolates. We assembled sequences for 27 new trichomonasvirus strains across all known TVV species, with 17 of these assemblies representing coding-complete genomes. Using a strategy of *de novo* sequence assembly followed by taxonomic classification, we discovered a fifth species of TVV that we term Trichomonas vaginalis virus 5 (TVV5). Six strains of TVV5 were assembled, including two coding-complete genomes. These TVV5 sequences exhibit high sequence identity to each other, but low identity to any strains of TVV1-4. Phylogenetic analysis corroborates the species-level designation. These results substantially increase the number of coding-complete TVV genome sequences and demonstrate the utility of mining publicly available transcriptomes for the discovery of RNA viruses in a critical human pathogen.

## INTRODUCTION

Trichomonas vaginalis viruses (TVVs) are monosegmented (i.e., nonsegmented) double-stranded RNA (dsRNA) viruses that infect the parasitic protozoan *Trichomonas vaginalis* (Goodman et al., 2011b). They constitute genus *Trichomonasvirus* in family *Totiviridae* (Goodman et al., 2011b) and are related to monosegmented dsRNA viruses that infect some other parasitic protozoa, namely, viruses that infect *Leishmania* spp. and constitute genus *Leishmaniavirus* in family *Totiviridae* (Tarr et al., 1988) and viruses that infect *Eimeria* spp. and constitute proposed genus *Eimeriavirus* in family *Totiviridae* (Wu et al., 2016). Monosegmented dsRNA viruses that infect *Giardia* spp. and constitute genus *Giardiavirus* (currently classified in family *Totiviridae*, though probably destined for separation) are more distantly related to trichomonasviruses (Wang and Wang, 1986).

The prevalence of *Trichomonas vaginalis* as a sexually transmitted human pathogen is a major reason for interest in the trichomonasviruses. *T. vaginalis* is an extracellular parasite that attaches to epithelial cells of the genitourinary tract of both men and women, and is the causative agent of trichomoniasis, the most common non-viral sexually transmitted infection worldwide. Trichomoniasis is associated with perturbed vaginal microbiota, premature delivery, low birth weight, infertility, and increased transmission and acquisition of other infectious diseases, including HIV-1 and HPV (Mercer and Johnson, 2018). Currently, the anti-microbial drug metronidazole is used to treat trichomoniasis, but drug resistance is rising, indicating a need for alternative therapies (Leitsch, 2016). Trichomonasviruses are known to increase the virulence of *T. vaginalis* by increasing the degree of inflammation during infection. Vaginocervical epithelial cells sense Trichomonasviruses via recognition of dsRNA by Toll-like receptor 3, which triggers a proinflammatory response through the NFkB and Interferon Regulating Factor-3 pathways (Fichrova et al., 2012). Increased inflammation is associated with premature delivery (Jun et al., 2000) and higher transmission and acquisition of infectious diseases, including HIV-1 (Alfano et al., 2002). This antiviral response has been demonstrated in response to TVV1. It is unknown whether this is a conserved response to all TVVs or rather is limited to a subset of these viruses.

A somewhat curious aspect of trichomonasviruses is that they comprise at least four different species (*Trichomonas vaginalis virus 1, Trichomonas vaginalis virus 2, Trichomonas vaginalis virus 3*, and *Trichomonas vaginalis virus 4*) (Bessarab et al., 2000, 2011; Goodman et al., 2011a; Tai and Ip, 1995) and that many *T. vaginalis* isolates are found to be simultaneously infected with different combinations of these species, including some isolates infected with all four (Goodman et al., 2011a; Rivera et al., 2017). This finding raises the question of the relative roles played by each species and how they may affect the virulence of *T. vaginalis*. It also presents the possibility of biologically relevant interactions among the different trichomonasvirus species, which may give rise to differential effects on the *T. vaginalis* host and ultimately the human superhost.

Transcriptome data deposited in public databases, such as sequence reads in the Sequence Read Archive (SRA) database and transcript assemblies in the Transcriptome Shotgun Assembly (TSA) database, both maintained at the National Center for Biotechnology Information (NCBI; Bethesda, MD, USA), are proving to be a boon for virus discovery (e.g., Nibert et al., 2018). For the current study, we decided to screen the public transcriptome data for *T. vaginalis* in an effort to discover additional trichomonasvirus strains. We found that the TSA database is devoid of transcript assemblies for *T. vaginalis* to date but that the SRA database contains sequence reads from a number of different transcriptome studies of this human pathogen. In the end from these SRA data sets, we were able to assemble complete or partial coding sequences for 27 new strains across all four currently recognized trichomonasvirus species. We supplemented this work by determining the complete coding sequences for two other new trichomonasvirus strains by de novo sequencing. Notably, we implemented *de novo* assembly methods and sensitive homology searches of distantly related sequences to discover a fifth species of TVV with two coding-complete gapless genomes and four partial assemblies. We term this novel trichomonasvirus TVV5. Comparisons of these new trichomonasvirus sequences enhance our understanding of several basic features of these viruses.

## RESULTS

### Screening for trichomonasvirus sequences in public transcriptome data

The Sequence Read Archive (SRA) database at NCBI includes accessions from several RNA-Seq transcriptome studies of *T. vaginalis*, deposited under five BioProjects from four institutions in the USA, Germany, and Korea (Bradic et al., 2017; Gould et al., 2013; Song et al., 2017; Woehle et al., 2014) (Table 1). These accessions encompass sequence reads from at least thirteen different *T. vaginalis* isolates. For this report, we screened these SRA data sets for the presence of sequence reads matching any of the four known species in genus *Trichomonasvirus* (Bessarab et al., 2000, 2011; Goodman et al., 2011a, 2011b; Tai and Ip, 1995). Briefly, we applied Discontiguous MegaBLAST at NCBI for performing the database searches, using three to six previously reported nt sequences for each trichomonasvirus species as queries. Multiple queries were used for each species in an effort to increase the numbers of identified hits from divergent strains of each species that might have been present in these isolates.

**Table 1.**
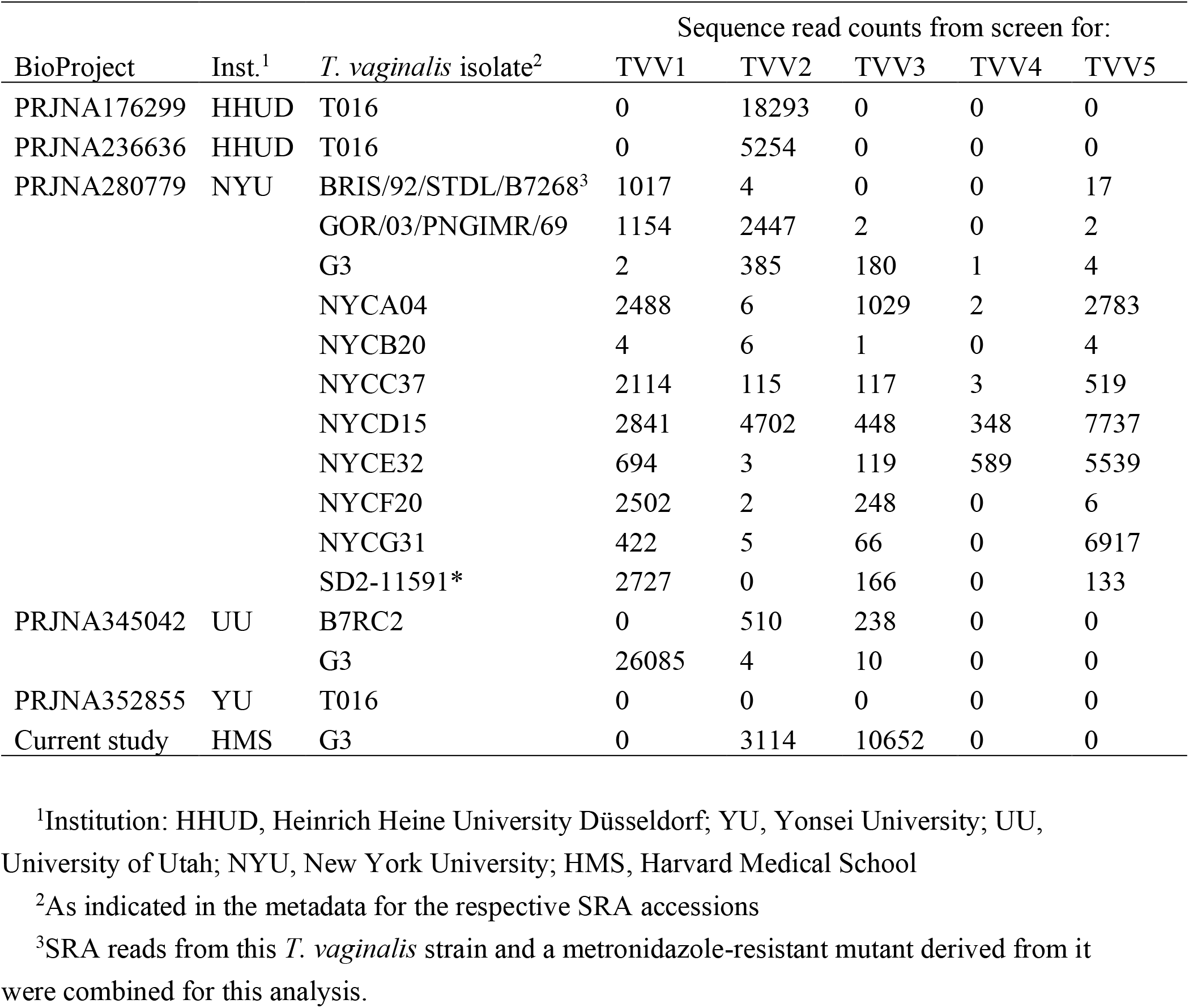
Screens of SRA data sets at NCBI for trichomonasvirus-matching sequence reads

Results of these screens are summarized in Table 1. Sequence reads matching strains of all four known trichomonasvirus species were found, often in large numbers. Only two of the sixteen SRA data sets that we distinguished for screening were concluded to be negative for strains of all four species (0 to 13 hits per species): the data set for *T. vaginalis* isolate T016 from BioProject PRJNA352855 and the data set for *T. vaginalis* isolate NYCB20 from BioProject PRJNA280779. The other fourteen data sets that we distinguished for screening were concluded to be positive for strains of at least one species each (67 to 26,085 hits per species): ten data sets positive for TVV1, seven data sets positive for TVV2 (including two from the same isolate and institution), nine data sets positive for TVV3, and two data sets positive for TVV4. Broken down by *T. vaginalis* isolate, isolates G3 (as reported from BioProject PRJNA345042, University of Utah (UU)) and BRIS/92/STDL/B7268 were positive for TVV1 only; isolate T016 (as reported from BioProjects PRJNA176299 and PRJNA236636, Heinrich Heine University Düsseldorf (HHUD)) was positive for TVV2 only; isolate GOR/03/PNGIMR/69 was positive for TVV1 and TVV2; isolates NYCA04, NYCF20, NYCG31, and SD2-11591* were positive for TVV1 and TVV3; isolates G3 (as reported from BioProject PRJNA345042, New York University (NYU)) and B7RC2 were positive for TVV2 and TVV3; isolate NYCC37 was positive for TVV1, TVV2, and TVV3; isolate NYCE32 was positive for TVV1, TVV3, and TVV4; and isolate NYCD15 was positive for all four species (Table 1). In total, the screening results suggest the identification of 27 new trichomonasvirus strains by this approach, as additionally examined below.

### Assembly of new trichomonasvirus sequences

Having coding-complete sequences for virus strains is useful for confirming expected features such as open reading frames (ORFs) and ribosomal frameshifting motifs, for identifying conserved features not previously recognized, for identifying phenotypically important sequence variations, and for allowing robust phylogenetic comparisons. We therefore next undertook to assemble the TVV-matching reads into coding-complete sequences for as many of the 27 suggested new trichomonasvirus strains as we could. For this effort, we used the programs CAP3 (Huang and Madan, 1999) and CLC Genomics Workbench to assemble the TVV-matching reads into contigs, and we also separately performed de novo assembly of contigs from the SRA data sets, using the program rnaSPAdes (Bushmanova et al., 2018), for corroboration of the results. Through this combination of approaches, we were able to generate and confirm coding-complete sequences for 13 new trichomonasvirus strains: ten TVV1 strains (TVV1-G3(UU), TVV1-BRIS/92/STDL/B7268, TVV1-GOR/03/PNGIMR/69, TVV1-NYCA04, TVV1-NYCC37, TVV1-NYCD15, TVV1-NYCE32, TVV1-NYCF20, TVV1-NYCG31, and TVV1-SD2-11591*) and three TVV2 strains (TVV2-T016, TVV2-GOR/03/PNGIMR/69, and TVV2-NYCD15). In addition, we were able to generate and confirm nearly coding-complete sequences for four other new trichomonasvirus strains: one other TVV2 strain (TVV2-B7RC2; small 3’ truncation) and three TVV3 strains (TVV3-B7RC2, TVV3-NYCA04, and TVV3-NYCD15; single small gap in each). Lastly, for the remaining ten trichomonasvirus strains indicated by the findings in Table 1 (two other TVV2 strains, six other TVV3 strains, and two TVV4 strains), we were able to generate and confirm three to six contigs of ≥300 nt in length for each, allowing them also to be included in the comparisons that follow. RPKM values, using the final full sets of reads used for generating the final assemblies (further details in Materials and Methods), ranged from 1.2 to 31 for the thirteen coding-complete assemblies (median, 5.2), 0.3 to 2.9 for the four nearly coding-complete assemblies (median, 0.8), and 0.4 to 1.5 for the longest contig from each of the ten partial assemblies (median, 0.6) (Table S1). Median sequencing depth values ranged from 10 to 621 for the thirteen coding-complete assemblies (median, 43), 6 to 16 for the four nearly coding-complete assemblies (median, 10), and 3 to 10 for the longest contig from each of the ten partial assemblies (median, 3) (Table S1).

### Discovery of a fifth species of Trichomonas vaginalis virus

Following our map-to-reference strategy to characterize additional strains of existing trichomonasviruses, we next devised an approach that would enable discovery of divergent viruses in these same isolates of *Trichomonas vaginalis*. A *de novo* assembly approach was implemented, whereby all adapter-trimmed RNA sequencing reads of a given isolate were assembled using De Bruijn graph-based methods. These assemblies were dynamically translated into the protein space and compared to the NBCI nonredundant ‘nr’ protein database using DIAMOND (Buchfink et al., 2015). Assemblies were compared to the top 1% of matching sequences in the reference database and a taxonomic origin was thus assigned. Assemblies assigned to viral taxa were retrieved and analyzed.

This assembly-first approach successfully reconstructed the virus sequences obtained from the previous map-to-reference strategy. It also enabled the search for virus strains sufficiently divergent from existing TVV sequences. Accordingly, isolate NYCE32 was found to carry a virus that could only confidently be assigned to genus *Trichomonasvirus* with no species-level determination. This 5042 bp assembly exceeded the length of any known TVV strain, though it shares the familiar TVV genomic architecture of two large overlapping reading frames of approximately equal length. Querying this assembly using nucleotide BLAST demonstrated a 42% match to its closest TVV sequence (TVV2-UR1) in the NCBI nonredundant nucleotide ‘nt’ database. As *Trichomonasvirus* species are demarcated by <50% sequence identity, this opened the possibility that NYCE32 harbored a novel TVV species.

To test whether additional Tvag isolates harbored similar sequences, we used the map-to-reference approach with the NYCE32 assembly to identify matching reads in any other sample. This ultimately led us to discover homologous sequences in five additional isolates (NYCA04, NYCC37, NYCD15, NYCG31, and SD2-11591*), resulting in six assemblies of a divergent *Trichomonasvirus*. Three of these assemblies are coding-complete, one assembly encodes a full-length CP protein, and the remaining two sequences are partial assemblies not fully spanning a viral gene. For the five sequences that cover the predicted CP/RdRp junction, a ribosomal frameshift site can be deduced. This heptanucleotide slippery sequence is GGGCCCC, which is the same motif used by TVV2. CP/RdRp fusion proteins determined from these six assemblies share >77% sequence identity with each other yet <50% sequence identity with any strain of TVV1-4.

Each *trichomonasvirus* species determined to date forms a distinct monophyletic clade. To examine the divergence of these six novel assemblies from currently established TVV species, we constructed a phylogenetic tree of CP/RdRp protein sequences from all newly determined transcriptome assemblies and NCBI TVV genomes (Fig. 1). These six assemblies were found to form a monophyletic clade with 100% branch support. This clade is positioned as a sister branch to TVV2, reflecting the next closest viral relative and a finding strengthened by the shared predicted ribosomal slippery sequence. Given the evidence presented thus far, namely a strong monophyletic clade, high within-group sequence identity and low sequence identity to TVV1-4, and a genome length that exceeds any known TVV strain, we contend that these six assemblies represent six strains of a novel *Trichomonasvirus*: TVV5.

**Figure 1.**
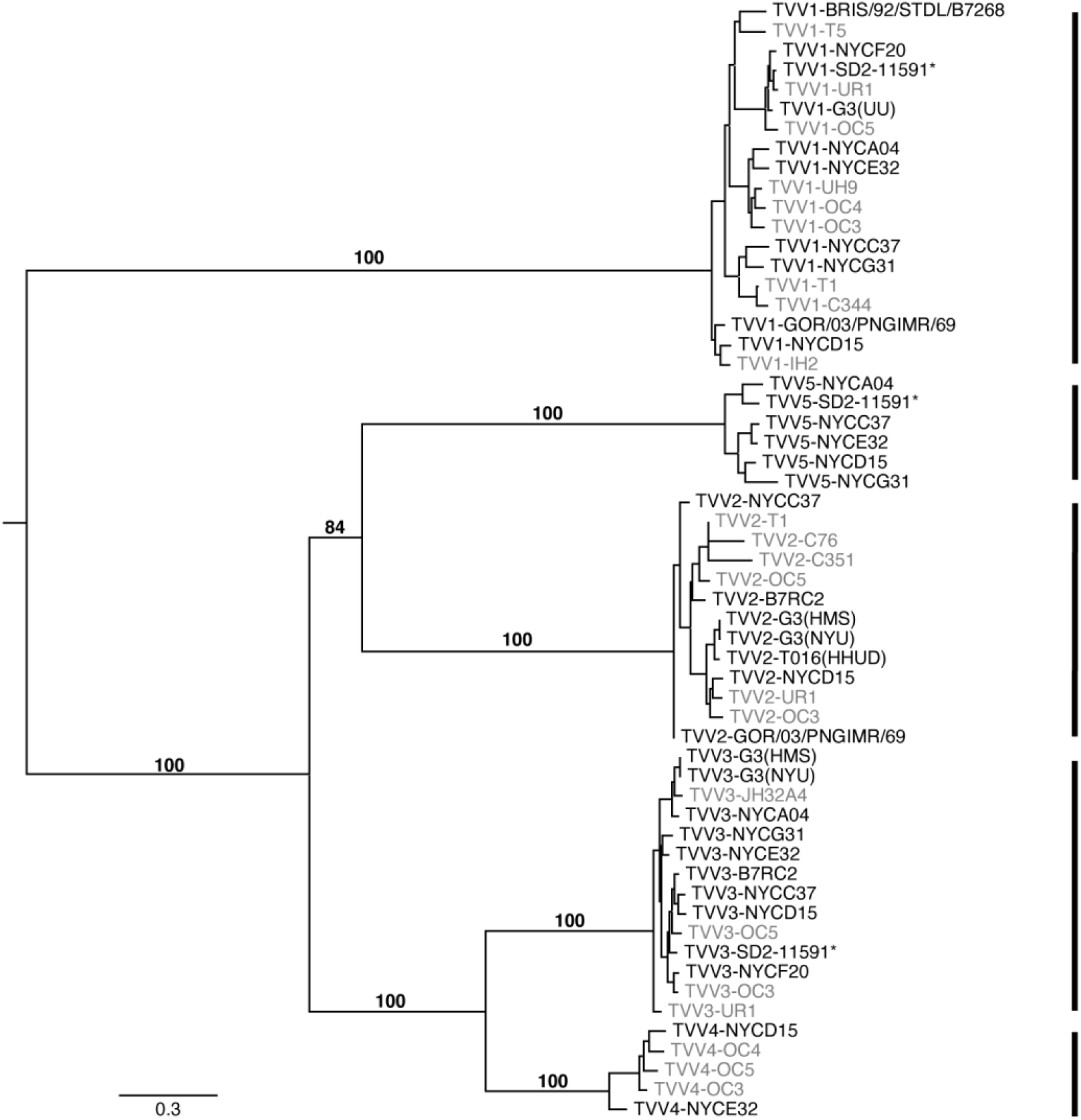
Phylogenetic tree of TVV1-5. CP/RdRp fusion protein sequences from determined from the novel assemblies presented in this study as well as reference TVV genomes in NCBI GenBank. New assemblies are labeled in black, and reference genomes are labeled in gray. All sequences were aligned using MAFFT (algorithm: L-ins-i). A phylogenetic tree was constructed using IQ-TREE on the LANL webserver (hiv.lanl.gov). The ‘choose best model and apply’ option was used to select the optimal substitution model. This was determined to be the model ‘JTT+F+I+G4’. 1000 ultrafast bootstraps were run. BOOSTER was used to optimize branch-level support values. These branch-level bootstrap values are shown in percentage values. The tree was visualized using FigTree.

### Reexamination of T. vaginalis isolate G3

Our results for the presence of trichomonasvirus sequences in the analyzed transcriptome data sets (Table 1) include inconsistent findings for *T. vaginalis* isolate G3. In particular, G3 from University of Utah (BioProject PRJNA345042) is positive for TVV1 only whereas G3 from New York University (BioProject PRJNA280779) is positive instead for TVV2 and TVV3. Since isolate G3 is readily obtainable from the American Type Culture Collection (ATCC; accession PRA-98) and also since G3 is the reference isolate for which the *T. vaginalis* whole genome sequence draft has been reported (Carlton et al., 2007), we newly obtained isolate G3 from the ATCC and examined it for the presence of trichomonasviruses.

For detecting TVV1, TVV2, and/or TVV3 strains that may be carried by isolate G3, we first designed primer pairs based on the TVV1-G3(UU), TVV2-G3(NYU), and TVV3-G3(NYU) sequences described above. RNA was then extracted directly from the cells of *T. vaginalis* isolate G3 that were present in the original sample sent from the ATCC, followed by RT–PCR using the respective primer pairs. In this manner, we consistently failed to obtain an amplicon of the expected size using the primer pair based on TVV1-G3(UU) but succeeded in obtaining amplicons of the expected sizes using the primer pairs based on TVV2-G3(NYU) or TVV3-G3(NYU). Moreover, when the latter amplicons were subjected to Sanger sequencing, we found them to be 100% identical in sequence to the respective region of TVV2-G3(NYU) (amplicon length excluding primers, 715 nt) and 99.6% identical in sequence to the respective region of TVV3-G3(NYU) (amplicon length excluding primers, 798 nt). Based on these results, we concluded that the trichomonasvirus results obtained for *T. vaginalis* isolate G3 reported from New York University are representative of those for *T. vaginalis* isolate G3 currently available from the ATCC.

Because the assembled sequences for TVV2-G3(NYU) and TVV3-G3(NYU) from BioProject PRJNA280779 are not coding complete, we also performed an RNA-Seq analysis of *T. vaginalis* isolate G3 that we obtained from the ATCC, in an effort to complete the coding sequences of these trichomonasvirus strains. Following enrichment of dsRNA from this isolate after limited passage in culture in our lab, the sample was submitted for RNA-Seq analysis by a commercial vendor (Quick Biology, Pasadena, CA). Results revealed the presence of many sequence reads matching TVV2 and TVV3, but not TVV1 or TVV4 (Table 1), consistent with the results from BioProject PRJNA280779 described above. Additionally, reads sufficient in number and coverage were obtained to assemble complete coding sequences for both TVV2-G3(HMS) and TVV3-G3(HMS) (HMS reflecting that this study was performed at Harvard Medical School; see Tables S1 and S2 for assembly statistics). These new assemblies exhibit 100% sequence identity with the amplicons for portions of these viruses described above and averages of 99.7% and 99.3% sequence identity, respectively, with the partial assemblies for TVV2-G3(NYU) and TVV3-G3(NYU).

### Comparisons of trichomonasvirus sequences

Untranslated regions (UTRs) at the termini of viral genomes are important sites for replication and/or viral packaging. Previous reports (Goodman et al. 2011, JVI) have found conserved elements across all strains of each TVV species. UTRs routinely exhibit more genomic flexibility than protein coding regions, as insertion-deletion events in the latter result in deleterious frameshifts and yield defective viral proteins. However, as noted above, UTRs can play vital functional roles as well which place constrains genetic variation at these sites. All TVVs exhibit sizable 5’-UTRs which could contain functional elements such as Internal Ribosomal Entry Sites (IRESes).

To search for functional elements in the UTRs, all novel TVV assemblies were compared to the currently available TVV sequences in the NCBI GenBank database. Per species, these sequences were aligned and any gaps (corresponding to insertion/deletions between strains) were plotted (Fig. 2). Most insertion-deletion events were found to occur in the UTRs, especially the 5’-UTR. In TVV1, TVV2, TVV3, and TVV5 there were distinct regions in the 5’-UTR that lacked any gaps. As TVV1 has the most sequences, the pattern can be most confidently seen with this virus. Two gap-free zones exist where no nucleotides have been inserted or deleted. To investigate whether genetic variation at these nucleotides is constrained, nucleotide conservations plots were generated for each TVV species (Fig. 3). The conservation plot for TVV1 shows two peaks of high sequence identity at the two gapless regions. The beginning of the CP gene is also highly conserved. These patterns of long 5-UTRs with distinct zones with constrained nucleotide variation and no insertions or deletions point to a functional role for the 5’-UTR of *Trichomonasviruses*. These observations allow for the possibility of a functional IRES in one or more TVV species.

**Figure 2.**
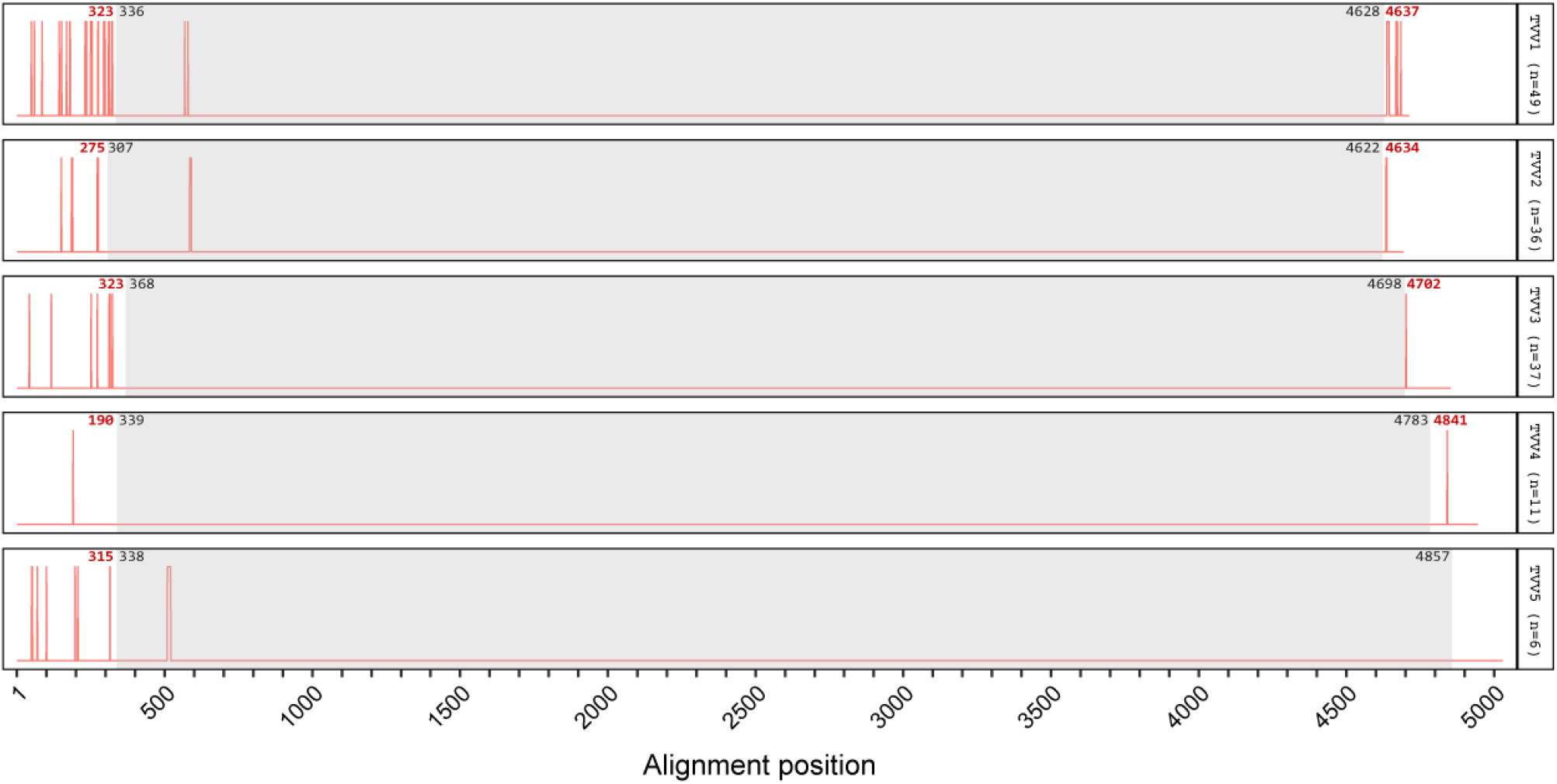
Gap plots delineating all insertion-deletion events across the genomes of TVV1-TVV5. Indels were found extensively in the genomic UTRs, but distinct gapless regions within the UTRs suggest conserved functional elements. Per TVV species, newly generated assemblies were combined with all coding-complete and partial sequences in NCBI Genbank and aligned using MAFFT L-ins-i. The unsequenced ends of partial sequences in the multiple sequence alignment were masked to prevent bias from missing residues. The alignment was analyzed with a custom script implemented in R. Sites with gaps are shown in red. The numbers in red indicate the nearest gap to the coding sequence boundaries, colored in black. The gray boxes denote the coding region of each virus species.

**Figure 3.**
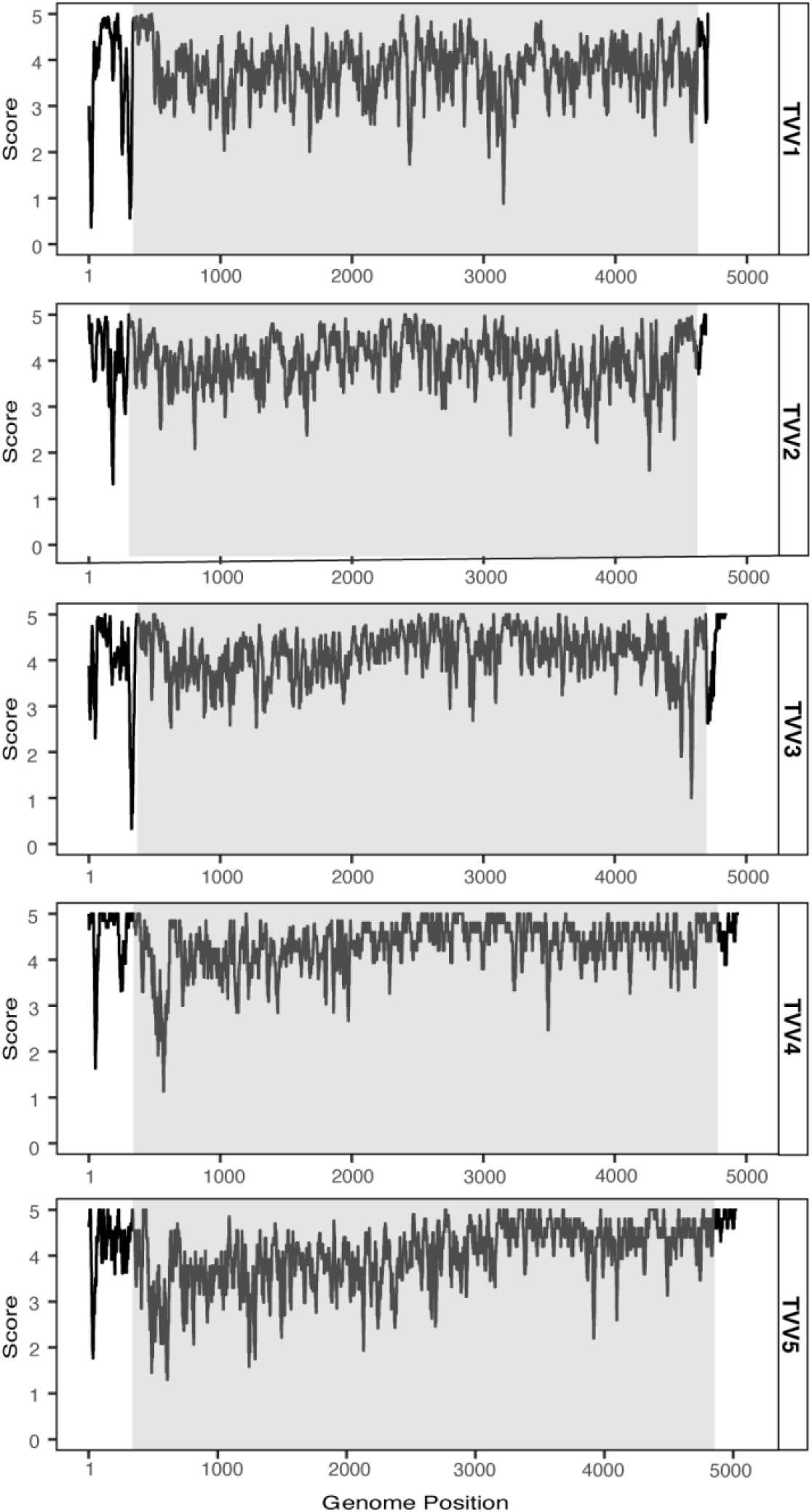
Nucleotide conservation plots for TVV1-TVV5. Partial and full-length sequences were retrieved from NCBI and combined with transcriptome derived full-length genomes per each species. This analysis was implemented in R. A sliding window of 15 nt was chosen for smoothing. EDNAFULL substitution matrix was used, in which a 5 denotes perfect identity at a given position. Gray box represents the coding sequence.

Insertions and deletions were found to be largely excluded from the coding regions of TVV genomes. No strains of TVV3 or TVV4 possessed any gaps within the coding sequence. All insertions/deletions were in the 5’- or 3’-UTRs. One strain of TVV5 (TVV5-NYCA04) possesses a 12 bp deletion near the beginning of the CP coding sequence, corresponding to a skip of 4 amino acids. This strain is also likely divergent from the rest, possessing an 18 amino acid N-terminal extension. TVV1 and TVV2 each had two GenBank sequences with apparent insertions within the CDS. The TVV1 strains are small amplicon sequences resulting from the same study on an Iranian patient cohort (Heidary et al, 2013). TVV1-SH8 (GenBank accession AB701566) is a 149 bp amplicon sequence and contains an inserted guanosine residue at position 568 relative to the whole genome alignment with all other TVV1 strains. TVV1-SH4 (GenBank accession AB701562) is a 142 bp amplicon sequence with an inserted adenosine residue at position 579. In both cases, the inserted nucleotide results in a frameshift that would yield a truncated capsid protein with a divergent C-terminus. Removal of the single nucleotide insertion restores the protein translation to perfect identity to other known TVV1 sequences. Both TVV2 sequences were from the same study on a Cuban patient cohort (Fraga et al, 2012). TVV2 strains C76 and C351 (GenBank accessions JF436870 and JF436871, respectively) each contain one multiple-nucleotide insertion near the beginning of the capsid gene. TVV2-C76 possesses the insertion ‘AAGAAA’ at positions 585-590, and TVV2-C351 possesses the insertion ‘TAA’ at positions 588-590. These insertions maintain the coding frame and would result in the introduction of one or two amino acids into the capsid protein.

TVV1 genomes were found to possess a conserved CUUUUUGCAC element in the 5’-UTR of full-length viruses. The sole exception is TVV1-NYCC37 which has an extra uracil residue resulting in CUUUUUUGCAC. This uracil-rich element is found in all full-length TVV1 genomes but not in any strains of TVV2, TVV3, TVV4, or TVV5. The 3’-UTRs of each species demonstrate fairly strong sequence conservation within a species, with little similarity across species. These species-specific UTR elements could play a role in segregation of viral components during the prevalent co-infections of multiple Trichomonasviruses in a single trichomonad.

Interestingly, our newly described TVV1 sequences are all coding-complete, have 5’-termini that match or exceed that of previously reported TVV1 genomes, and 3’-termini within 20 nt of reference TVV1 genomes. Three of the seven transcriptome-derived TVV2 sequences are coding-complete, with a fourth sequence nearly coding-complete. Three of nine TVV3 sequences are coding-complete. Neither of the two novel TVV4 sequences is coding-complete. Genome completeness was found to be correlated with sequencing depth (Figure S1). To evaluate whether number of mapped reads is related to TVV species, a nonparametric Kruskal-Wallis test was conducted to evaluate whether each TVV species had equivalent numbers of mapped reads per sample (normalized as RPKM values). RPKM values were found to vary significantly across species (p = 0.029), presenting the possibility of different levels of each TVV within a host protozoan (Fig. S2).

To evaluate sequence conservation across TVV species, CP/RdRp amino acid sequences were determined from newly assembled sequences and coding-complete sequences deposited in NCBI. These sequences were globally aligned per species using CLUSTAL Omega. Pairwise identity matrices for all strains in a given TVV species are provided in Supplementary Tables S3-S7.

All TVV1 genomes assembled in the course of this study are coding-complete, allowing for the most detailed analysis of any *Trichomonasvirus* species. Almost all the TVV1 CP/RdRp sequences are 1430 aa in length; TVV1-UR1-1 and SD2-11591* both contain an MGIP N-terminal extension, bringing their coding sequence lengths to 1434 aa. TVV1 sequences share a triple serine carboxy terminus except for TVV1-NYCC37 which encodes an STS at the C-terminus. Average sequence identity across the TVV1 CP/RdRp sequences is 85.9%. Scanning over the CP/RdRp sequence with a 10 amino acid window reveals a single area of the genome with <50% amino acid conservation. This region occurs halfway through the CP/RdRp sequences, centered on amino acid 701 (705 for TVV1-UR1-1 and SD2-11591*). Interestingly, absolute conservation is observed twenty residues upstream, the site of the ribosomal frameshift.

For TVV2, the average sequence identity for the fusion protein is 88.3%. The coding-complete TVV2 CP/RdRp sequences are 1438 aa. The N-terminus of these TVV2 sequences contains a MASTL motif, except for the CP/RdRp from TVV2-T016-HHUD which encodes a MAATL motif. The C-terminus of all full-length TVV2 fusion proteins ends in PVYV. TVV3 CP/RdRp proteins share an average sequence identity of 92.1%. The N-terminus of TVV3 begins with MSAPEPLNTEVR, and all TVV3 fusion protein C-termini end with a GHGLRSG motif. Analysis of the TVV4 protein sequences was impeded by the fact that no novel coding-complete TVV4 genomes could be constructed from the SRA data. The TVV4-NYCD15 and TVV4-NYCE32 assemblies contain partial capsid coding sequences. Protein sequences derived from these sequences have a pairwise identity score of 90.3%. These sequences were aligned to the CP coding sequences of the three TVV4 genomes in GenBank: TVV4-OC3, TVV4-OC4, and TVV4-OC5. TVV4-NYCD15 shares the MSAI N-terminal motif with TVV4-OC3 through TVV-OC5. Neither new assembly reached the CP C-terminus.

The TVV5 CP/RdRp proved to be the largest of all *Trichomonasvirus* polymerase proteins. The three complete or nearly complete assemblies encode a fusion protein of 1506 aa. Examination of all TVV5 strains shows a largely conserved N-terminus. The exception is TVV5-NYCA04 which possesses a guanosine at genome position 283 instead of the adenosine found in all other sequences with coverage at that position. This alters the codon from AUA to AUG. Along with a nearby single nucleotide deletion, these elements allow for an earlier predicted initiation of translation, resulting in an 18 amino acid N-terminal extension of the CP and CP/RdRp proteins for this strain. Three TVV5 sequences maintain coverage through the CP/RdRp C-terminus. In all three sequences, the protein ends in PAVPIAT.

Programmed ribosomal frameshifts are instrumental in TVV biology, giving rise to the catalytic CP/RdRp protein. All strains sequenced in this report demonstrate the absolute conservation of each species’ heptanucleotide frameshift motif (TVV1: CCUUUUU, TVV2/TVV5: GGGCCCC, TVV3/TVV4: GGGCCCU). No variation at these sites was observed for any strain of any species.

One common observation across all TVV species is that while the amino acids surrounding the ribosomal frameshift are strictly conserved, 10-30 amino acids downstream exhibit extreme divergence. For example, the TVV2 ribosomal frameshift site occurs at residue 699Q. TVV2 positions 693-702 are perfectly conserved across all strains. However, just downstream is a region of low conservation, dropping below 50% at positions 737 and 738 after which modest sequence identity (80-95%) is seen again. The evolutionary constraint of the ribosomal frameshift may cause higher diversifying selection pressure in the neighboring genomic space.

To evaluate any conserved RNA structures across trichomonasviruses, secondary structure analysis was conducted on all TVVs. The first and last 100 nucleotides from TVV1-OC3, TVV2-OC3, TVV3-OC3, TVV4-OC3, and TVV5-NYCE32 were analyzed for secondary RNA structure at the 5’ or 3’ end of the coding strands of their genomes using RNAfold (Lorenz et al., 2011). As expected from previous reports (Goodman et al. 2011), a conserved 5’ stem-loop was again found in the 5’ terminus of all Trichomonasviruses. More interestingly, TVV2, TVV3, TVV4, and TVV5 were found to possess a conserved double stem-loop structure at their 3’ genomic terminus (Fig. 4). This structure consists of two adjoining stem-loops with no interspersing nucleotides. These structures extend to within 5 nucleotides of the 3’ terminus in TVV2, TVV3, and TVV5. The TVV4 double stem-loop structure was found 21 nucleotides upstream of the 3’ terminus. This conserved RNA feature could play a role in packaging the genome into virions and/or recognition by the viral RNA-dependent RNA polymerase.

**Figure 4.**
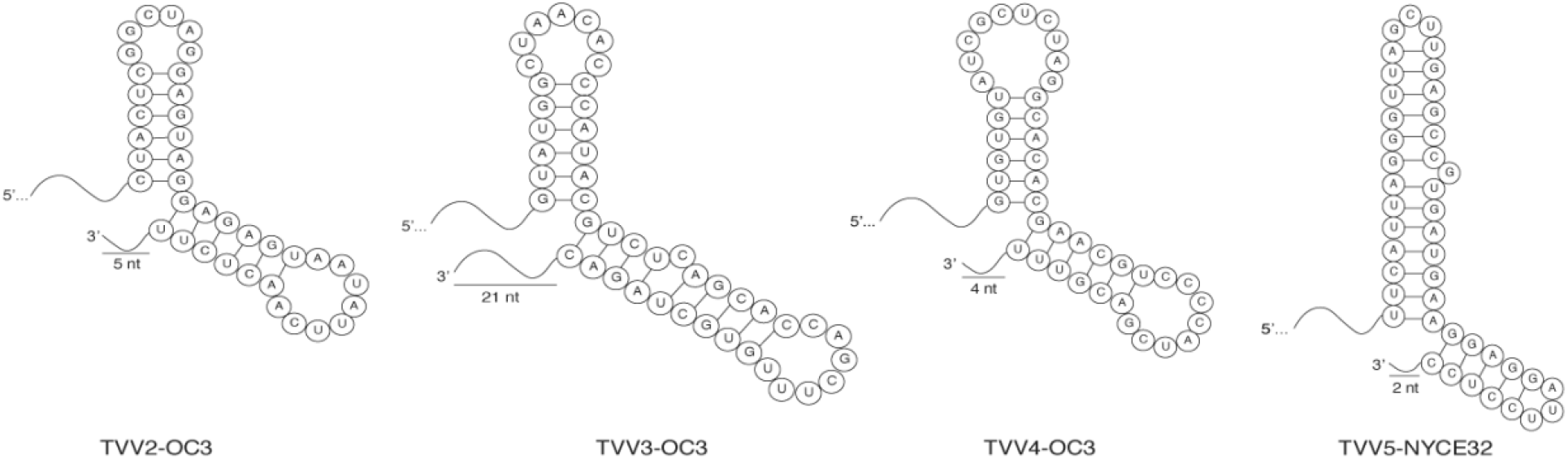
Conserved double stem-loop structure in the 3’ genomic terminus of TVV2-TVV5. Secondary structure analysis identified a conserved feature in TVV2-OC3, TVV3-OC3, TVV4-OC3, and TVV5-NYCE32. This feature is found within 21 nucleotides of the 3’ terminus of the coding strand of each virus and consists of two adjoining stem-loops with no interspersing nucleotides.

## DISCUSSION

Discovery of a fifth species of TVV underscores the utility of mining viral sequences from publicly available transcriptomes. A map-reads-to-reference strategy was useful in characterizing additional strains of known viral species. An orthogonal ‘assemble-first, classify-later’ approach ultimately allowed us to uncover a new virus infecting the human pathogen *Trichomonas vaginalis*. These complimentary methods enable a robust exploration of sequence data while incorporating the flexibility needed for finding new viruses. Each virus assembly held up to rigorous scrutiny, corroborated in all cases by both reference-based and *de novo* assembly methods. Table S1 shows the normalized read count per virus assembly presented in this report, expressed as reads per kilobase per million reads (RPKM). Most assemblies have an RPKM of <5. Relatedly, a majority of these assemblies have less than 20x coverage (Table S2). These values are strong enough to allow assembly of viral sequences and resulting in the assembly of several coding-complete genomes presented in this report. However, this might also explain the number of partial assemblies. For the newly assembled TVV4 sequences, both of which are partial assemblies, each RPKM was <1 and median coverage was <10x. Viral sequencing mining was demonstrably successful using these untargeted datasets, yet methods for enriching viruses in RNA-seq library preparation may be considered for reliable determination of coding-complete viral genomes.

Goodman et al. (2011a) reported a conserved, short 5’-terminal region of sequences (36 or 37 nt) in strains TVV1-UH9, TVV1-UR1, TVV1-OC3, TVV1-OC4, and TVV1-OC5 (GenBank accessions HQ607516.1, HQ607513.1, HQ607516.1, HQ607521.1, and HQ607523.1), which was missing from previously reported sequences for strains TVV1-1, TVV1-T5, and TVV1-IH2 (GenBank accessions U08999.1, U57898.1, and DQ270032.1) (Kim et al., 2007; Tai and Ip, 1995; Su and Tai, 1996) and is also missing from the subsequently reported sequence for strain TVV1-C344 (Fraga et al., 2012). This short sequence extension is noteworthy because it allows formation of a long 5’-terminal stem–loop structure, which seems likely to be involved in RNA stability and/or other functions in TVV1 replication. All of the new TVV1 sequences reported here extend into this conserved 5’-terminal region, providing further evidence that the sequences for TVV1-1, TVV1-T5, TVV1-IH2, and TVV1-C344 are likely 5’-truncated. The fact that the new TVV1 sequences reported here appear to be less but still partially truncated themselves is not surprising given that RNA-seq-derived transcript assemblies, in our experience, are often missing a few terminal residues at each end of respective transcripts.

Strains of all five trichomonasvirus species have long 5’ untranslated regions (UTRs), >280 nt each, as well as numerous AUG codons preceding the first in-frame AUG codon in ORF1 (Goodman et al., 2011). These features, also evident in the new strains reported here, suggest to us that at least some of these 5’ UTR sequences contribute to forming an internal ribosome entry site/structure (IRES) required for trichomonasvirus translation, as shown or suggested for several other members of family *Totiviridae* or related dsRNA viruses (Chiba et al., 2018; Garlapati and Wang, 2005). The increased number of complete coding sequences now available for alignment as a consequence of the current study additionally reveals a previously unemphasized fact, namely, that indels between strains are found in the 5’ UTR of each trichomonasvirus species, as shown in Fig. 2. These indels may mark sequence locations that are not directly involved in mediating essential functions at the RNA level or in forming essential RNA secondary structures. Indels between strains are also found in the 3’UTR of each trichomonasvirus species, but are generally not found in the long coding region that occupies most of the genome.

With regard to trichomonasvirus strains in *T. vaginalis* isolate G3, our de novo sequencing results and those from the study by Bradic et al. (2017) based at New York University (BioProject PRJNA280779) concur in identifying TVV2 and TVV3, but not TVV1 or TVV4, in this isolate. Moreover, the high levels of nucleotide-sequence identity (≥99.3%) between TVV2-G3(HMS) and TVV2-G3(NYU) and between TVV3-G3(HMS) and TVV3-G3(NYU) are consistent with limited divergence of these viruses while the *T. vaginalis* isolate was cultured at the ATCC and/or then separately at the two recipient institutions. On the other hand, the discrepancies found for the viruses in this isolate from the study at University of Utah (BioProject PRJNA345042) (presence of TVV1, but not TVV2, TVV3, or TVV4) are harder to explain, especially given that this isolate was again obtained from the ATCC according to BioSample metadata from that study. One possibility would seem to be that the SRA data for isolates G3 and B7RC2 from the University of Utah study might have been transposed in the database, since the current study identified both TVV2 and TVV3 in isolate B7RC2, as expected instead for isolate G3 based on other results. However, TVV2-B7RC2 and TVV3-B7RC2 are substantially divergent from TVV2-G3(HMS) and TVV3-G3(HMS) (≤85.4% nt-sequence identity), which makes this explanation seem unlikely to us. At this stage, then, these discrepancies remain unexplained, but they highlight the need for investigators to confirm the virus content of *T. vaginalis* isolates used in each new study that is focused on these viruses.

The current study also identifies discrepancies in the trichomonasvirus content of *T. vaginalis* isolate T016. From two studies at Heinrich Heine University Düsseldorf (BioProjects PRJNA176299 and PRJNA236636) (Gould et al., 2013; Woehle et al., 2014), isolate T016 is found here to contain TVV2, but not TVV1, TVV3, or TVV4, and the TVV2 sequences derived from those two studies are found to be identical. In contrast, isolate T016 from Yonsei University (BioProject PRJNA352855) (Song et al., 2017) is found here to be negative for all four trichomonasvirus spp. In this case, since the discrepancies involve a single trichomonasvirus species, the *T. vaginalis* isolate T016 from Yonsei University might have simply been “cured” of TVV2 during culture, as has been reported to occur in some other cases (Wang et al., 1987; Men-Fang et al., 1993).

The double stem-loop structure found in TVV2-TVV5 is a striking feature, possibly playing a role in the viral lifecycle. Conserved secondary structures have been found in other RNA viruses to be involved in replication and virion formation. While this double stem-loop structure varies in length somewhat between the viruses, the maintenance of its overall shape in TVV2-TVV5 suggests a functional role in the TVV lifecycle. However, that this feature was not found in TVV1 is not entirely surprising. The evolutionary distance is considerable between TVV1 and the rest of the TVV species, as deduced from phylogeny as well as facets of viral biology such as translation strategies. TVV2, TVV3, TVV4, and TVV5 seem to employ a −1 ribosomal frameshifting mechanism for translation of the CP/RdRp fusion protein while TVV1 uses a −2 ribosomal frameshifting strategy. TVV1 may possess its own characteristic secondary structures to fulfill vital roles of its lifecycle. Future biochemical studies dissecting the functional potential of TVV RNA species appear especially warranted.

## MATERIALS AND METHODS

### Screens

Transcriptome datasets in the NCBI Sequence Read Archive (SRA) database were screened for *Trichomonasvirus*-matching sequencing reads. This analysis was carried out with a locally implemented BLAST (Camacho et al., 2008) instance to retrieve hits followed by a de-deduplication step to exclude erroneous reads cross-mapping to other TVV species. Discontiguous megablast (‘stand_alone_blast.sh’) was first run with multiple TVV species-specific queries against the respective SRA datasets using an e-value threshold of 1e-9. Blastn (‘cleanup_blast.sh’) was subsequently run on those hits against a local blast database containing 80 TVV sequence accessions retrieved from GenBank using an e-value threshold of 10. This confirmed how many of the initial hits indeed best matched the queried TVV species. Table 1 shows these results for the number of TVV reads for each respective species in each dataset. Normalized coverage values, expressed as RPKM, were calculated for each trichomonasvirus assembly, shown in Table S1. Sequencing depth was also calculated for each trichomonasvirus assembly by mapping each set of TVV reads to a reference strain for that species using BWA mem (Li H., 2013). The samtools (Li H. et al., 2013) ‘depth’ function was used to determine number of mapped reads per nucleotide position, and a median was calculated per virus assembly using the ‘median’ function in the statistical programming language, R (R Core Team, 2013). These coverage values are presented in Supplemental Table S2.

### High-throughput sequencing

*T. vaginalis* isolate G3 was purchased from the American Type Culture Collection (ATCC ID: PRA-98). This isolate was minimally cultured in Diamond’s Modified Medium (Fouts & Kraus, 1980) for three passages at late-exponential phase. Total RNA was isolated using TRIzol. Contaminating ssRNA was depleted with a 2M LiCl incubation at −20°C overnight. Final enrichment was achieved with a nuclease digestion using DNase I and S1 nucleases (Promega Corporation, Madison, WI). Enriched dsRNA was shipped overnight on dry ice to Quick Biology (Pasadena, CA) for sequencing. The dsRNA-seq library was prepared according to KAPA KK8540 RNA HyperPrep kit with 201-300 bp insert size (KAPA Biosystems, Wilmington, MA) using 25-50 ng of total dsRNA as input. Final library quality and quantity were analyzed by Agilent Bioanalyzer 2100 and Life Technologies Qubit 3.0 Fluorometer. Paired-end 150 bp reads were sequenced on an Illumina HiSeq 4000 (Illumina Inc., San Diego, CA). Sequences were demultiplexed to remove barcodes at the sequencing facility.

### Bioinformatic analysis

Adapters and other technical sequences were trimmed from the paired-end reads using TrimGalore v0.6.3 (https://github.com/FelixKrueger/TrimGalore) with the following parameters: paired, stringency=5, quality=20. Bacteriophage ΦX174 positive-control spike-in reads were depleted using ‘BWA-mem’ with default parameters. Reads were assigned to their respective Trichomonasvirus using the BLAST workflow above (i.e., initial mapping with ‘discontiguous megablast’ followed by a cleanup step with ‘blastn’). After binning the reads, species-specific reads were *de novo* assembled into a draft genome using CAP3 (parameters: -o 21 -p 66 -s 300 -z 2). In addition to this map-to-reference approach, *de novo* assembly strategy was also taken using rnaSPAdes v3.13.0 on all G3(HMS) reads for draft viral genome assembly sensitive to any significant structural variations. The draft assembly for each virus was refined by mapping the reads to the assembly and generating a 50% consensus sequence using a mapping script (‘refine_contigs.sh’) that implements BWA-mem, samtools, and vcftools (Danecek et al., 2011). Both approaches converged on the same result, yielding coding complete genomes from *T. vaginalis* isolate G3 for TVV2 and TVV3. This *de novo* assembly approach was likewise used for discovery of divergent *Trichomonasvirus* sequences. Identification of divergent sequences was performed using DIAMOND against the NCBI nonredundant (nr) protein database. The DIAMOND parameter ‘--top 1’ was used to force the lowest common ancestor (LCA) algorithm to consider the 99^th^ percentile of reference sequences to determine the taxonomic origin of each assembly. A custom Python script (‘diamondToTaxonomy.py’) was used to convert NCBI taxonomy IDs to full taxonomic lineages using the JGI-DOE taxonomy server (https://taxonomy.jgi.doe.gov).

### Phylogenetic tree construction

CP/RdRp fusion protein sequences were from determined from newly transcriptome-assembled TVV1-5 sequences and reference TVV genomes in NCBI. These sequences were aligned with MAFFT using the ‘L-ins-i’ algorithm (Katoh et al., 2005). A maximum likelihood phylogenetic tree was built from this alignment using IQ-tree (Nguyen et al., 2015) on the LANL webserver (hiv.lanl.gov). The ‘choose best model and apply’ option (Kalyaanamoorthy et al., 2017) determined that the model ‘JTT+F+I+G4’ was optimal and was thus used to build the tree. 1000 ultrafast bootstraps were conducted using UFBoot2 (Hoang et al. 2018), and a bootstrap tree was also created. The resulting maximum likelihood consensus tree and bootstrap tree was used for input into BOOSTER (Lemoine et al., 2018) implemented on the Pasteur Institute webserver (booster.pasteur.fr). Resulting phylogenetic tree was visualized using FigTree v1.4.4. Optimized bootstrap values are shown as branch-level support values in Fig. 1.

### Graphics

All plots were made in R using ggplot2 (Wickham, 2016). with extensive use of the tidyverse framework (Wickham et al., 2019). Figures were finalized in Adobe Illustrator.

### Data and code availability

RNA-sequencing reads were deposited to the NCBI Sequence Read Archive (SRA) database under accession SRX8785706. Bioinformatics code used for analysis in this study has been made freely available in the ‘TVV Transcriptome Mining’ repository at www.github.com/austinreidmanny/tvv-transcriptome-mining.

## ACKNOWLEDGMENTS

We wish to acknowledge and thank those scientists who made this study possible by depositing their sequence reads in the public databases. This research was funded in part by NIH Grant T32 AI007245 to the Ph.D. program in Virology at Harvard University (A.R.M., C.A.H.) and NIH Grant R01 AI132445 (A.R.M., C.A.H., M.L.N).

---

**Table S1.**
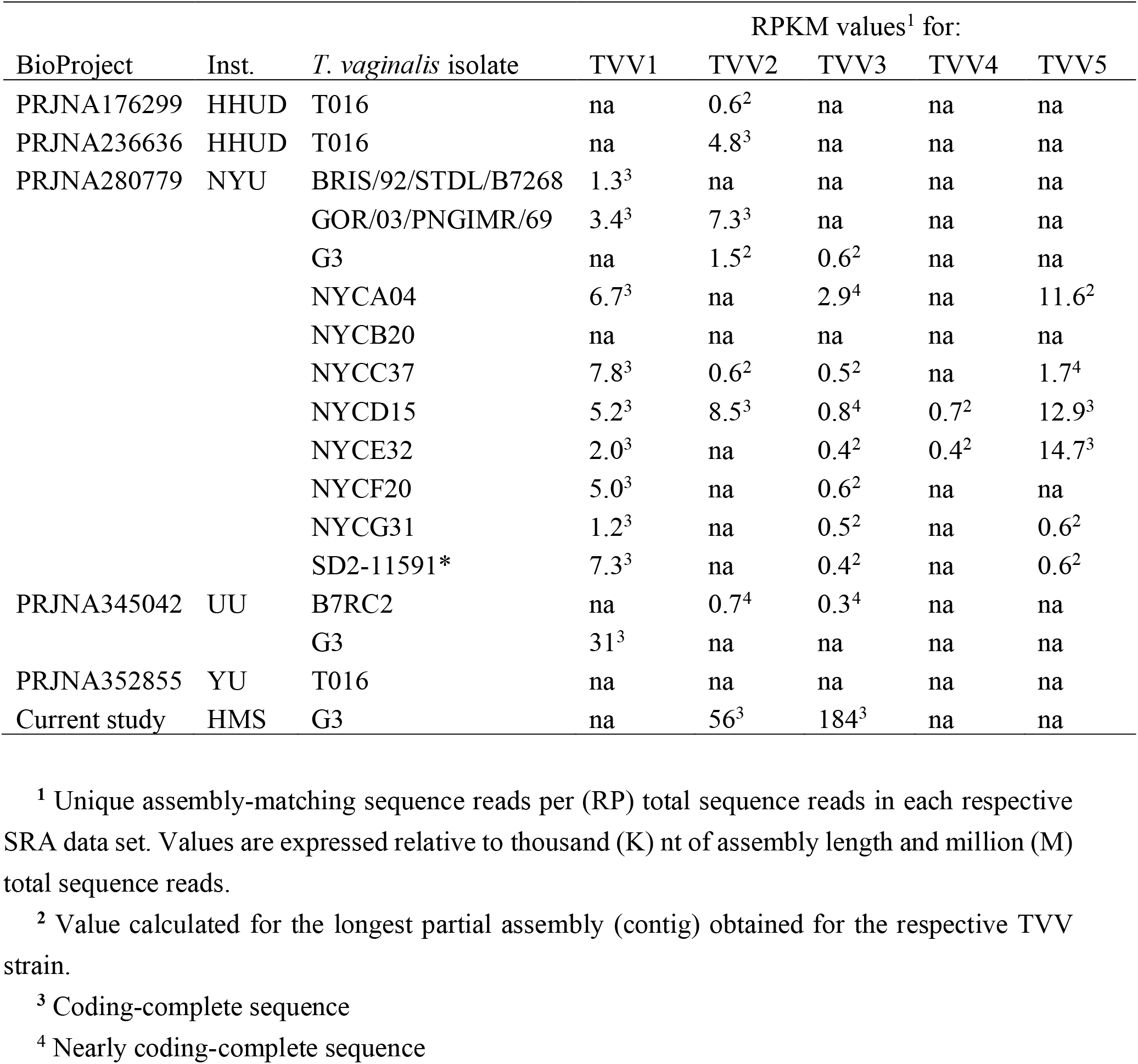
Sequence read counts for the new trichomonasvirus assemblies, expressed as RPKM values

| BioProject | Inst. | *T. vaginalis* isolate | RPKM values<sup>1</sup> for: |  |  |  |  |
| --- | --- | --- | --- | --- | --- | --- | --- |
|  |  |  | TVV1 | TVV2 | TVV3 | TVV4 | TVV5 |
| PRJNA176299 | HHUD | T016 | na | 0.6<sup>2</sup> | na | na | na |
| PRJNA236636 | HHUD | T016 | na | 4.8<sup>3</sup> | na | na | na |
| PRJNA280779 | NYU | BRIS/92/STD/L/B7268 | 1.3<sup>3</sup> | na | na | na | na |
|  |  | GOR/03/PNGIMR/69 | 3.4<sup>3</sup> | 7.3<sup>3</sup> | na | na | na |
|  |  | G3 | na | 1.5<sup>2</sup> | 0.6<sup>2</sup> | na | na |
|  |  | NYCA04 | 6.7<sup>3</sup> | na | 2.9<sup>4</sup> | na | 11.6<sup>2</sup> |
|  |  | NYCB20 | na | na | na | na | na |
|  |  | NYCC37 | 7.8<sup>3</sup> | 0.6<sup>2</sup> | 0.5<sup>2</sup> | na | 1.7<sup>4</sup> |
|  |  | NYCD15 | 5.2<sup>3</sup> | 8.5<sup>3</sup> | 0.8<sup>4</sup> | 0.7<sup>2</sup> | 12.9<sup>3</sup> |
|  |  | NYCE32 | 2.0<sup>3</sup> | na | 0.4<sup>2</sup> | 0.4<sup>2</sup> | 14.7<sup>3</sup> |
|  |  | NYCF20 | 5.0<sup>3</sup> | na | 0.6<sup>2</sup> | na | na |
|  |  | NYCG31 | 1.2<sup>3</sup> | na | 0.5<sup>2</sup> | na | 0.6<sup>2</sup> |
|  |  | SD2-11591* | 7.3<sup>3</sup> | na | 0.4<sup>2</sup> | na | 0.6<sup>2</sup> |
| PRJNA345042 | UU | B7RC2 | na | 0.7<sup>4</sup> | 0.3<sup>4</sup> | na | na |
|  |  | G3 | 31<sup>3</sup> | na | na | na | na |
| PRJNA352855 | YU | T016 | na | na | na | na | na |
| Current study | HMS | G3 | na | 56<sup>3</sup> | 184<sup>3</sup> | na | na |

<sup>1</sup> Unique assembly-matching sequence reads per (RP) total sequence reads in each respective SRA data set. Values are expressed relative to thousand (K) nt of assembly length and million (M) total sequence reads.
<sup>2</sup> Value calculated for the longest partial assembly (contig) obtained for the respective TVV strain.
<sup>3</sup> Coding-complete sequence
<sup>4</sup> Nearly coding-complete sequence

**Table S2.** Sequencing depth values for the new trichomonasvirus assemblies

| BioProject | Inst. | *T. vaginalis* isolate | Median sequencing depth values<sup>1</sup> for: |  |  |  |  |
| --- | --- | --- | --- | --- | --- | --- | --- |
|  |  |  | TVV1 | TVV2 | TVV3 | TVV4 | TVV5 |
| PRJNA176299 | HHUD | T016 | na | 7<sup>2</sup> | na | na | na |
| PRJNA236636 | HHUD | T016 | na | 102<sup>3</sup> | na | na | na |
| PRJNA280779 | NYU | BRIS/92/STD/L/B7268 | 19<sup>3</sup> | na | na | na | na |
|  |  | GOR/03/PNGIMR/69 | 20<sup>3</sup> | 39<sup>3</sup> | na | na | na |
|  |  | G3 | na | 10<sup>2</sup> | 4<sup>2</sup> | na | na |
|  |  | NYCA04 | 38<sup>3</sup> | na | 16<sup>4</sup> | na | 12<sup>2</sup> |
|  |  | NYCB20 | na | na | na | na | na |
|  |  | NYCC37 | 43<sup>3</sup> | 3<sup>2</sup> | 3<sup>2</sup> | na | 7<sup>4</sup> |
|  |  | NYCD15 | 48<sup>3</sup> | 68<sup>3</sup> | 7<sup>4</sup> | 7<sup>2</sup> | 14<sup>3</sup> |
|  |  | NYCE32 | 13<sup>3</sup> | na | 3<sup>2</sup> | 3<sup>2</sup> | 14<sup>3</sup> |
|  |  | NYCF20 | 48<sup>3</sup> | na | 6<sup>2</sup> | na | na |
|  |  | NYCG31 | 10<sup>3</sup> | na | 3<sup>2</sup> | na | 4<sup>2</sup> |
|  |  | SD2-11591* | 49<sup>3</sup> | na | 3<sup>2</sup> | na | 3<sup>2</sup> |
| PRJNA345042 | UU | B7RC2 | na | 13<sup>4</sup> | 6<sup>4</sup> | na | na |
|  |  | G3 | 621<sup>3</sup> | na | na | na | na |
| PRJNA352855 | YU | T016 | na | na | na | na | na |
| Current study | HMS | G3 | na | 34<sup>3</sup> | 161<sup>3</sup> | na | na |

<sup>1</sup> Unique assembly-matching sequence reads overlapping each nt position in the assembly.<sup>2</sup> Value calculated for the longest partial assembly obtained for the respective TVV strain.<sup>3</sup> Coding-complete sequence<sup>4</sup> Nearly coding-complete sequence

**Tables S3-S7 are provided as alignment matrices in the supplementary XLSX file.**

**Table S3**

Pairwise identity matrix of TVV1 CP/RdRp amino acid sequences, globally aligned using CLUSTAL Omega on the EBI-EMBL webserver. This includes the CP/RdRp protein sequence from novel assemblies as well as all coding-complete TVV1 genomes in the NCBI GenBank database.

**Table S4**

Pairwise identity matrix of TVV2 CP/RdRp amino acid sequences, globally aligned using CLUSTAL Omega on the EBI-EMBL webserver. This includes the CP/RdRp protein sequence from novel assemblies as well as all coding-complete TVV2 genomes in the NCBI GenBank database. Partial assemblies generated in this study that do not fully span the CP/RdRp protein are included.

**Table S5**

Pairwise identity matrix of TVV3 CP/RdRp amino acid sequences, globally aligned using CLUSTAL Omega on the EBI-EMBL webserver. This includes the CP/RdRp protein sequence from novel assemblies as well as all coding-complete TVV3 genomes in the NCBI GenBank database. Partial assemblies generated in this study that do not fully span the CP/RdRp protein are included.

**Table S6**

Pairwise identity matrix of TVV4 CP/RdRp amino acid sequences, globally aligned using CLUSTAL Omega on the EBI-EMBL webserver. Only partial assemblies not fully spanning the CP/RdRp protein were able to be generated for TVV4. These are aligned with the CP/RdRp protein from all coding-complete TVV4 genomes in the NCBI GenBank database.

**Table S7**

Pairwise identity matrix of TVV5 CP/RdRp amino acid sequences, globally aligned using CLUSTAL Omega on the EBI-EMBL webserver. The six novel assemblies generated in this study were used for this alignment.

**Figure S1.**
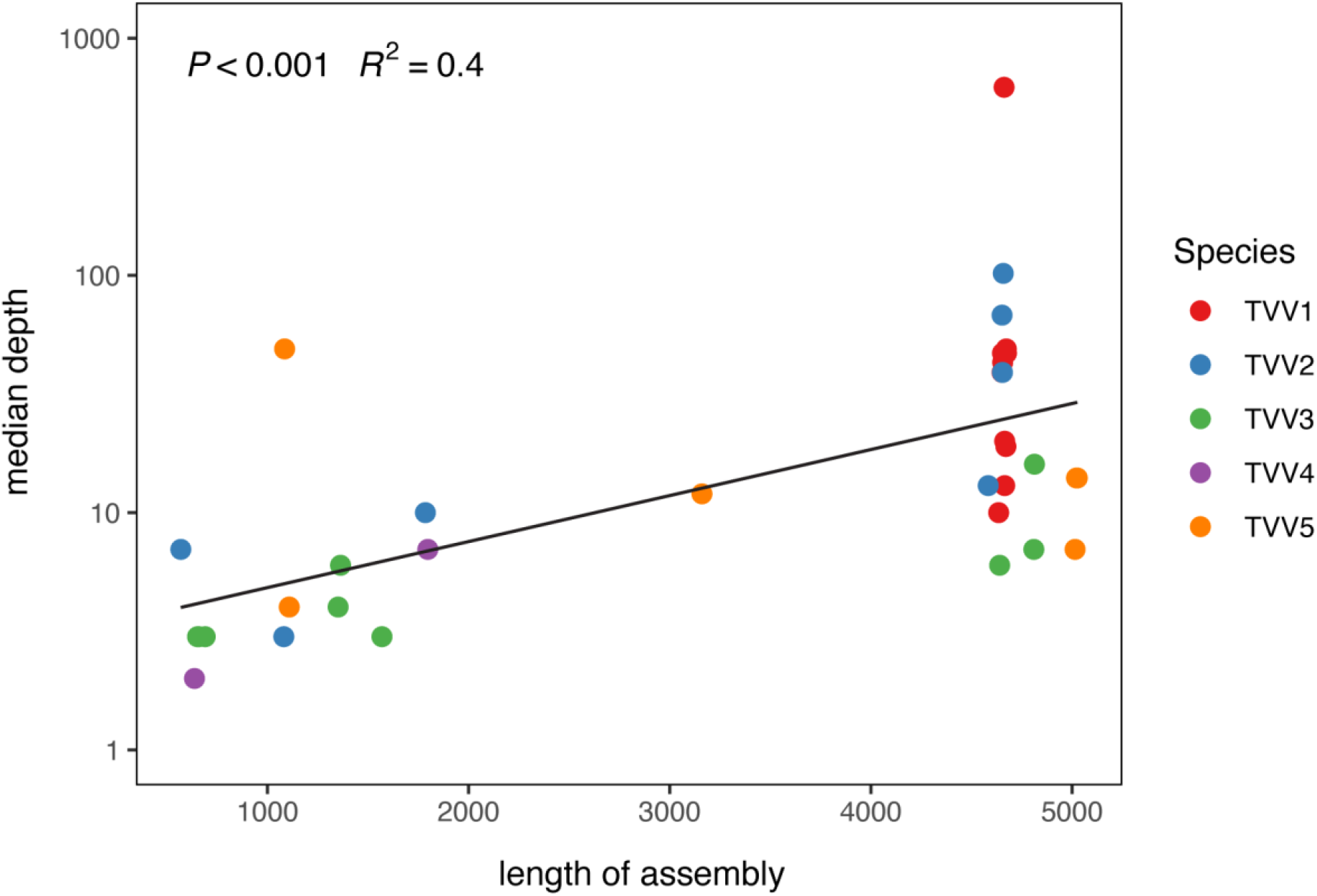
Distribution of TVV assembly lengths versus the median coverage depth across each sequence. For this dataset, obtaining a mapping depth of 10 reads per nucleotide increased the success rate of determining a coding-complete TVV genome from a metatranscriptome not enriched for viruses.

**Figure S2.**
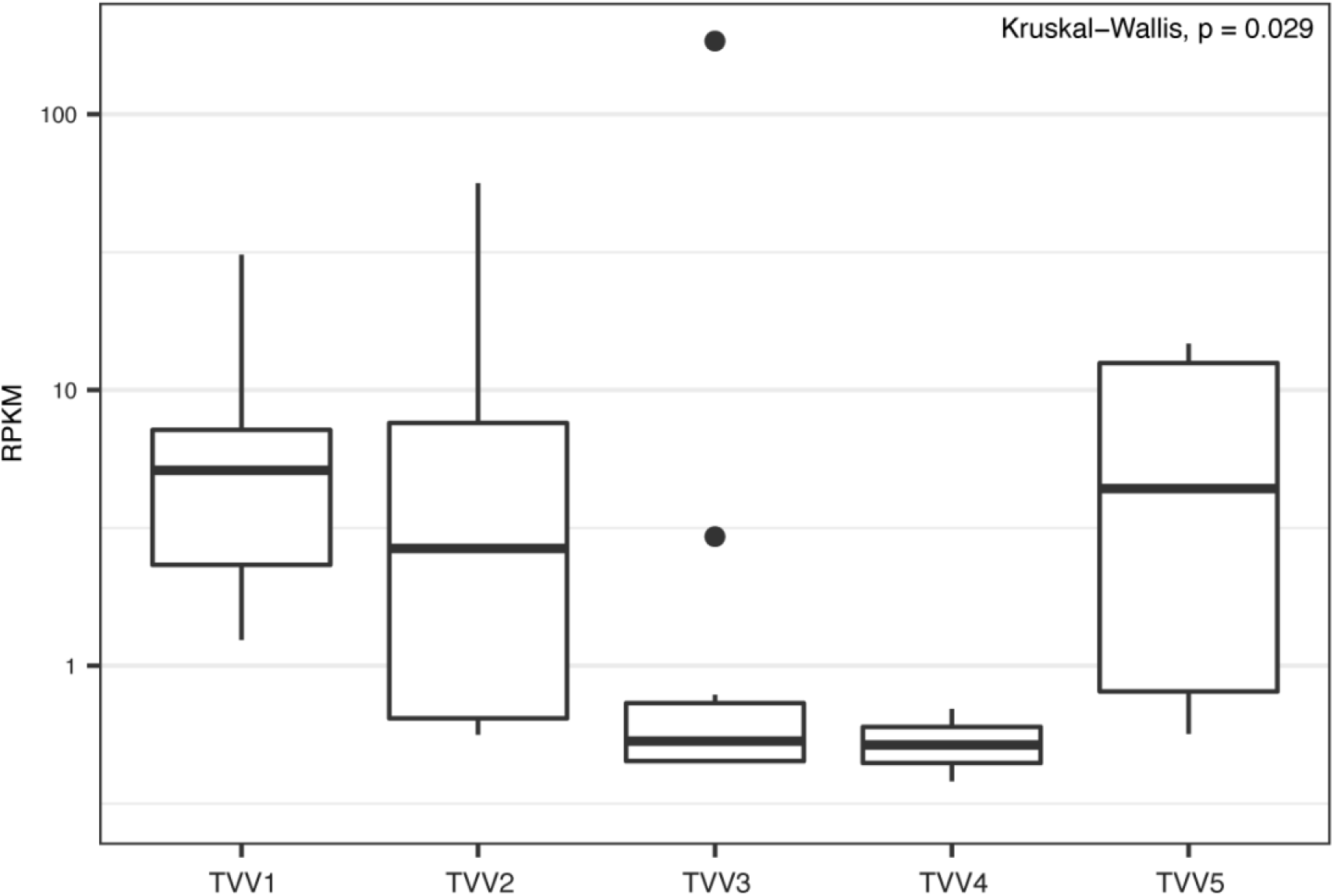
Mapped reads (normalized by RPKM) per species of TVV. The number of TVV reads varies across species, presenting the possibility of different levels of viral replication across TVVs.

